# Highly Mutable Linker Regions Regulate HIV-1 Rev Function and Stability

**DOI:** 10.1101/424259

**Authors:** Bhargavi Jayaraman, Jason D Fernandes, Shumin Yang, Cynthia Smith, Alan D Frankel

## Abstract

The HIV-1 protein Rev is an essential viral regulatory protein that facilitates the nuclear export of intron-containing viral mRNAs. Its sequence is organized into short, structured, functionally well-characterized motifs joined by less understood linker regions. We recently carried out a competitive deep mutational scanning study, which determined the relative fitness of every amino acid at every position of Rev in replicating viruses. This study confirmed many known constraints in Rev’s established interaction motifs, but also identified positions of mutational plasticity within these regions as well as in surrounding linker regions. Here, we probe the mutational limits of these linkers by designing and testing the activities of multiple truncation and mass substitution mutations. We find that these regions possess previously unknown structural, functional or regulatory roles, not apparent from systematic point mutational approaches. Specifically, the N- and C-termini of Rev contribute to protein stability; mutations in a turn that connects the two main helices of Rev have different effects in nuclear export assays and viral replication assays; and a linker region which connects the second helix of Rev to its nuclear export sequence has structural requirements for function. Thus, we find that Rev function extends beyond its characterized motifs, and is in fact further tuned by determinants within seemingly plastic portions of its sequence. At the same time, Rev’s ability to tolerate many of these massive truncations and substitutions illustrates the overall mutational and functional robustness inherent in this viral protein.

**Author Summary (non-technical summary):** HIV-1 Rev is an essential viral protein that controls a critical step in the HIV life cycle. It is responsible for transporting viral mRNA messages from the nucleus to the cytoplasm where they can contribute to the formation new virus particles. In order to understand how different regions of the Rev protein sequence are involved in its function, we introduced truncations and mass substitution mutations in the protein sequence and tested their effect on protein function. Through this study, we not only confirmed previous work highlighting known functionally important regions in Rev, but also found that a large portion of Rev, with little known functional roles influence Rev function and stability. We also show that although protein sequence is critical to its function, Rev can tolerate large variations to its sequence without disrupting its function significantly.

## Introduction

Proteins balance optimal functionality with mutational tolerance in order to adapt to changes in selection pressures [1]. Proteins with high tolerance for mutation are considered genetically “robust” or “plastic”, while proteins with a low mutability are considered genetically “fragile” or “brittle” [2]. The existence of a protein sequence as fragile or robust is a result of evolutionary pressures acting on the protein, such as mutation rate and biological function. Fragility and robustness are generally expected to couple with protein structure and disorder: disordered regions are expected to be robust while structured regions are comparatively brittle [3, 4].

Viral proteins maintain their function amidst many forces such as high viral polymerase error rates, immune pressures, and even competing selective pressures from overlapping reading frames and RNA structures. HIV-1 Rev is one such essential viral protein which facilitates the nuclear export of intron-containing viral RNAs that encode essential viral structural and enzymatic proteins and provide full length genomes for encapsidation [5, 6]. Rev experiences a high mutation rate in HIV (∼10^−3^ mutations per base per cell [7]) and is also overlapped with two other essential viral proteins, Tat and Env (Figure 1A). Overlap between viral genes is common [8], but most HIV-1 genes have at least one region with no overlap in which they can encode critical functional domains in an unconstrained way (as we recently observed with HIV-1 *tat* [9]). In contrast, *rev* shares coding information with other viral genes throughout its length (the only HIV-1 gene encoded in such a manner) meaning that its evolution is coupled to these overlapping genes.

**FIGURE 1:**
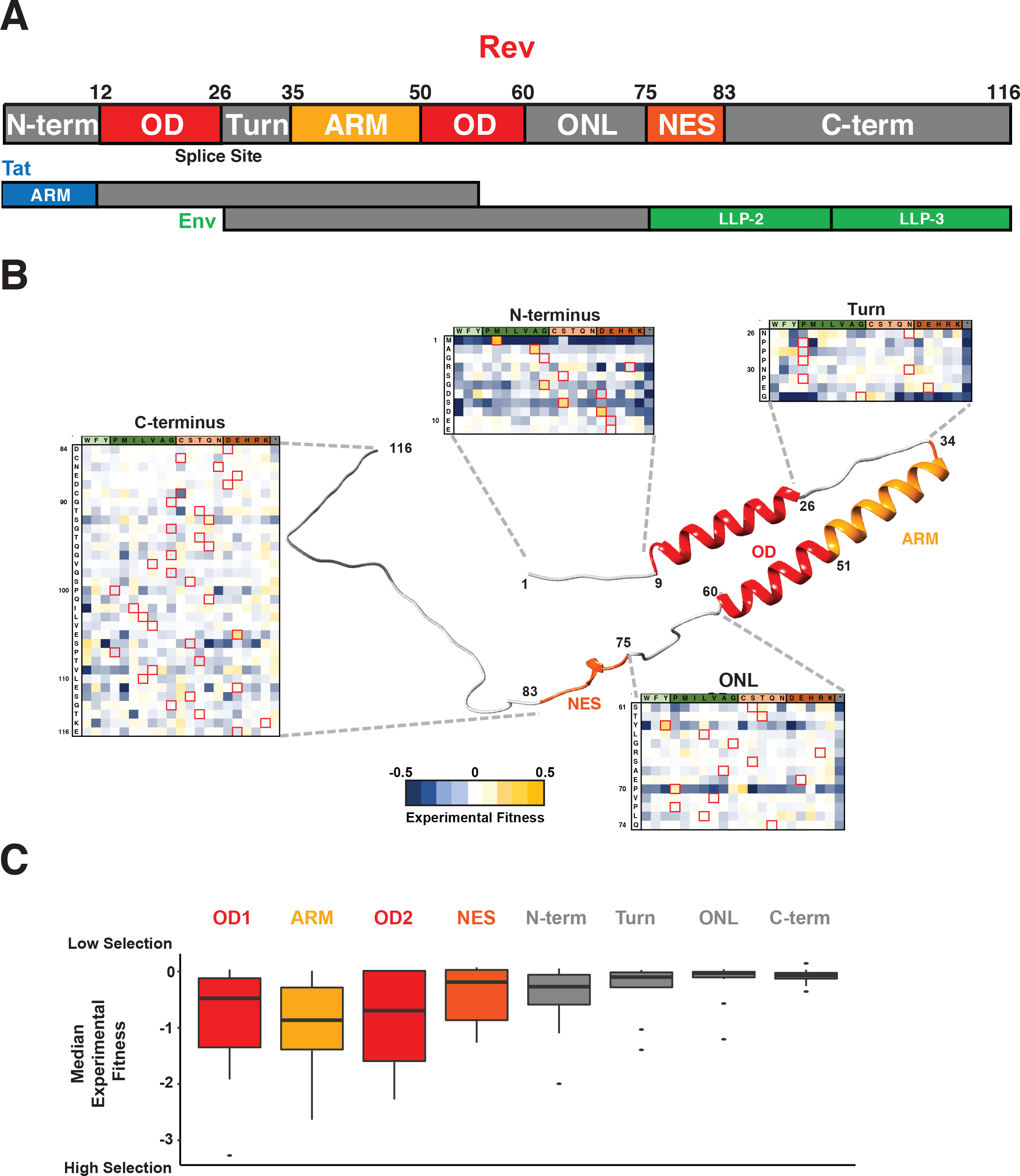
Structure and Organization of HIV-1 Rev. A) Domain organization of Rev protein (NL4-3/HXB2 numbering) with interaction surfaces in shades of red (OD: Oligomerization domain; ARM: Arginine-Rich Motif; NES: Nuclear Export Sequence) and linker regions in grey (N-term: N-terminus, Turn, ONL: OD-NES Linker). In the viral genome, the Rev coding sequence consists of two exons (Figure 1A; “splice site” denotes exon-exon junction as it would occur in the mRNA) both of which are contained entirely within (alternative reading frame) exons of two other HIV genes, Tat and Env. Putative structured domains of Tat (ARM: Arginine Rich Motif) and Env (LLP: Lentivirus lytic peptide) are shown along with their overlap with Rev domains. B) Structural model of Rev built using Rev crystal structures (residues 8-62 from PDBID: 3LPH; residues 76-83 from PDBID: 3NBZ; all other regions were built in PyMOL for visualization purposes). Also shown is the experimental fitness of every residue in the Rev linker regions (from [9]) C). Distribution of median experimental fitness values of each residue for different regions of Rev classified by domain organization (data from [9]). Specifically, there are 116 total data points each representing a single residue of Rev, each binned into the appropriate domain; the representative value for each residue is the median fitness of all 21 side chains (and the stop codon) at that position)

Because of the overlap, the true functional importance of each Rev residue is difficult to assess from traditional sequence conservation analyses and instead requires careful mutational dissection in non-overlapped contexts. To address this, we recently performed competitive deep mutational scanning (CDMS) in non-overlapped viral replication assays [9]. These experiments allowed us to examine for the first time, the amino acid preferences/fitness in Rev at each position, when unconstrained by overlapping genes.

Results from these experiments showed that, in general, the known structured/functional regions of Rev experience selective pressure for specific side chains while the linker regions between/flanking these structured domains appear to be relatively “plastic”, freely mutating between many different side chains with a variety of chemical properties (Figure 1B, C). However, the true extent of this plasticity is unclear, as the approach only examined single point mutations within a reference viral strain background.

In this study, we expand our analyses of the point mutational datasets to evaluate the limits of the structural and mutational plasticity of Rev’s structured domains and unstructured linker regions by designing deletion and mass substitution mutations and testing them for function in reporter assays and viral replication assays. Our results indicate that most of the linker regions possess previously unappreciated structural, functional or regulatory roles, not apparent from systematic point mutational approaches, while the structured regions tolerate more variation than expected. The N- and C-termini control Rev stability; mutations in a turn that connects the two main helices of Rev have different effects in nuclear export assays and viral replication assays, suggesting an alternate functional role besides nuclear export for this region; and a linker region connecting the second helix of Rev to its nuclear export sequence needs to be structured as a helix for function. Overall, our study illustrates for the first time how these linker regions are closely involved in Rev function either by direct functional roles or by more subtle regulatory roles. In sum, these experiments make evident Rev’s extraordinary genetic robustness, which is achieved via the use of malleable macromolecular interaction surfaces spaced by even more malleable linker regions that can more finely tune those interactions.

## Results

### Competitive deep mutational scanning (CDMS) data recapitulates Rev domain structure

Rev plays an essential role in the viral life cycle, coordinating the nuclear export of unspliced or partially spliced mRNAs for translation of key viral proteins or encapsidation of full-length viral genomes. Rev recognizes a ∼350-nt structured RNA within the intron known as the Rev Response Element (RRE) using an arginine-rich motif (ARM) for binding, and forms an oligomeric assembly on the RRE using two hydrophobic oligomerization motifs or domains (ODs). This Rev-RRE complex then recruits the host nuclear export factor Crm1, using a nuclear export sequence (NES) in Rev, together with RanGTP to transport these RNAs to the cytoplasm and thereby circumvent splicing [5, 6].

The three main functional regions of Rev (ARM, OD, and NES) are flanked by linker regions [10–14] (Figure 1A, B). The structure of the OD is formed by the interaction of two OD helices, where the second helix extends beyond the 15 residues of the helical ARM [12]. The NES forms its structure upon binding to Crm1 [15]. The regions that flank these functional domains include: a short N-terminal tail (N-term), a proline-rich turn between the first and second helices (turn), a linker between the second helix and the NES (OD-NES Linker: ONL) and a long, disordered C-terminal tail (C-term) [6, 16] (Figure 1B).

We utilized the fitness values for every possible Rev allele derived from a competitive deep mutational scanning (CDMS) dataset, where Rev was engineered into non-overlapped viruses and all substitutions at individual positions were competitively replicated, to deduce functional signatures of amino acid selection [9]. These data are consistent with our current biochemical and structural understanding of Rev, as we observe that the structured regions display low mutational tolerance (low median fitness in these regions indicates that only a few side chains are fit at each position) whereas the linker regions display high mutational tolerance (near-neutral median fitness indicates that almost all side chains are fit) (Figure 1B, C). The mutational plasticity of the linker regions is also apparent in the heat maps of all individual point mutations, further corroborating the underlying structural and functional boundaries (Figure 1B).

### The Rev OD is highly plastic at the sequence and structural levels

We first examined the amino acid requirements of the OD in more detail. It has long been known that Rev function requires binding of a homo-oligomer of 6-10 subunits to the RRE [16, 17]. The OD is comprised of two helices which contain hydrophobic residues necessary both for oligomerization and for interacting with host proteins, such as hnRNPs and RNA helicases [18–21]. The OD forms three hydrophobic surfaces with some residues shared between the surfaces: 1) a monomer stabilization interface in which side chains from each helix of the OD (residues 19, 22, 52 and 59) stabilize the monomer structure of Rev, 2) a dimer interface which supports formation of a Rev dimer upon nucleation of Rev assembly on the RRE (residues 18, 22, 55 and 59 and residue 21 in one dimer configuration), and 3) a higher-order interface opposite the dimer interface which allows the formation of the larger Rev oligomer (residues 12, 16 and 60) [10, 11, 22, 23] (Figure 2A).

**FIGURE 2:**
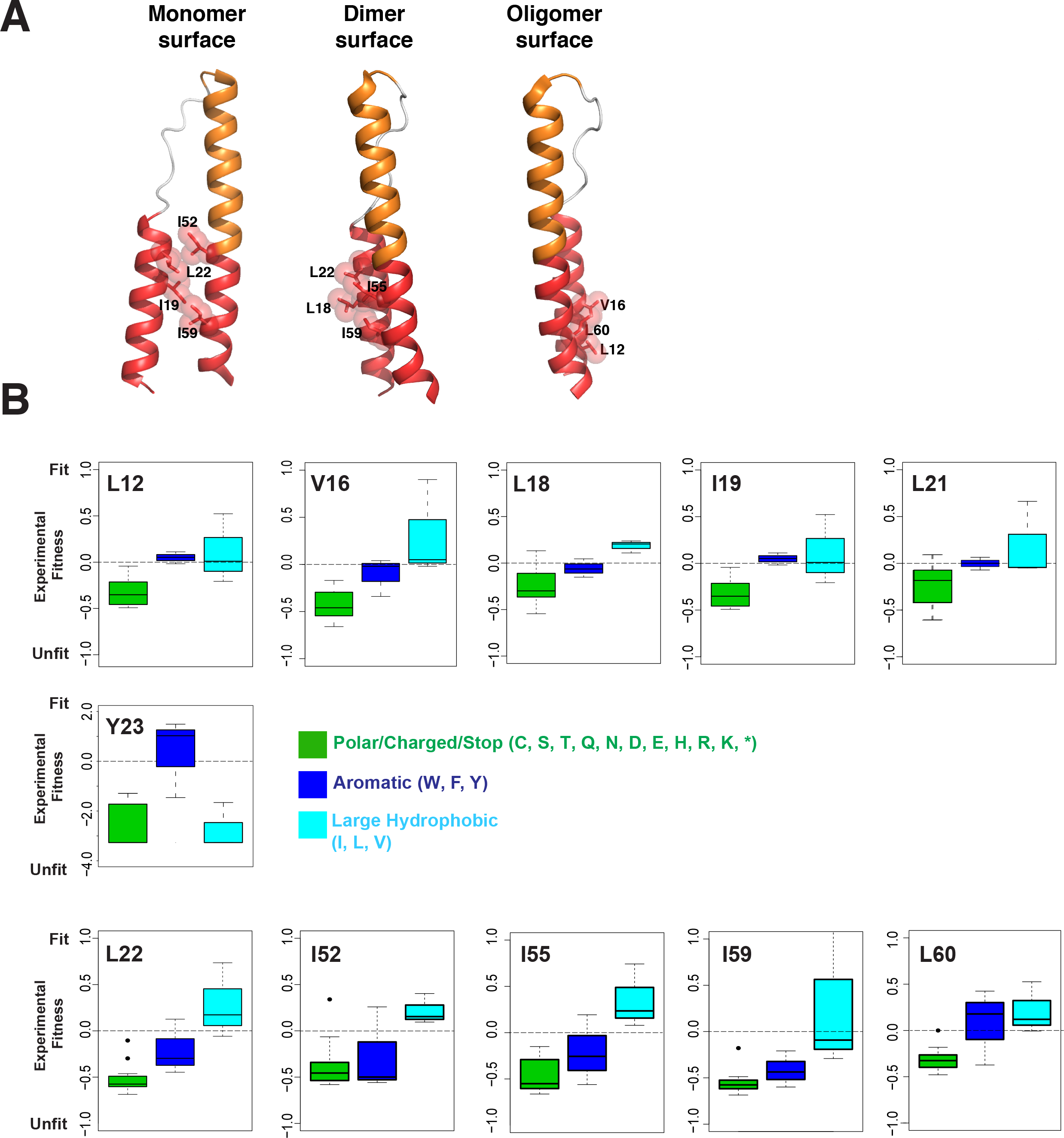
Selection of hydrophobic residues in the CDMS dataset [9] for the Rev Oligomerization Domain (OD) and nuclear export sequence (NES). A) Residues from the Rev OD involved in monomer, dimer or oligomer interface packing. Structures from PDBID 3LPH and 2X7L were used to generate the images. B) Amino acid preferences for important Rev OD contact residues using CDMS fitness values [9]. All residues display selection against polar/charged (C,S,T,Q,N,D, E, H, R, K) residues (and stop codons) (green) while some prefer large, hydrophobic side chains (teal) over aromatics (blue). Note change of scaling for Y23.

One very provocative feature of the Rev OD is that variation in the packing of its hydrophobic residues leads to multiple configurations of Rev dimers and oligomers [12, 24, 25]. This structural plasticity underlies the ability of Rev to assemble into a oligomeric complex on the RRE and raises the question of how constrained these side chains must be. CDMS data [9] show that interaction residues in each interface display a general preference for hydrophobicity and size with occasional preference for aliphatic side chains over aromatic ones (Figure 2B). For instance, in the dimer interface, residues 22, 55, and 59 prefer large aliphatic side chains while aromatic substitutions are equally as fit as aliphatic ones at residue 19. Similar patterns are seen in the higher-order oligomerization interface with residues 12 and 60 accepting any large hydrophobic, residues 16, 55 and 60 having distinct preferences for aliphatic residues. Interestingly, residue 23, although not part of currently known oligomeric interface configurations, strongly prefers F or Y (Figure 2B). In summary, this ability of the OD to make functionally neutral mutations between chemically similar side chains (e.g. I, L & V and in some cases W, F, and Y) at such functional critical positions increases the mutational robustness and prevents the “brittleness” that essential interfaces in other proteins can possess.

### The Rev NES and ARM balance fragility and plasticity

The Rev ARM serves as both the RRE-binding domain as well as a nuclear localization signal (NLS) through interactions with Importin-β [26]. It can recognize diverse RNA sites depending on the RNA structure and the orientation/placement of ARM residues [12, 17]. Additionally, residues in the ARM also enhance the monomer core structure through hydrogen bonds [10]. Given these numerous functional constraints, most residues of the ARM, including the non-basic residues Q36 and W45, show strong selection in the CDMS study (Figure 3A). Another residue, Q51, implicated in stabilizing an RNA-bound Rev dimer through hydrogen bonds [12] also shows strong selection, reaffirming its importance in Rev-RRE assembly.

**FIGURE 3:**
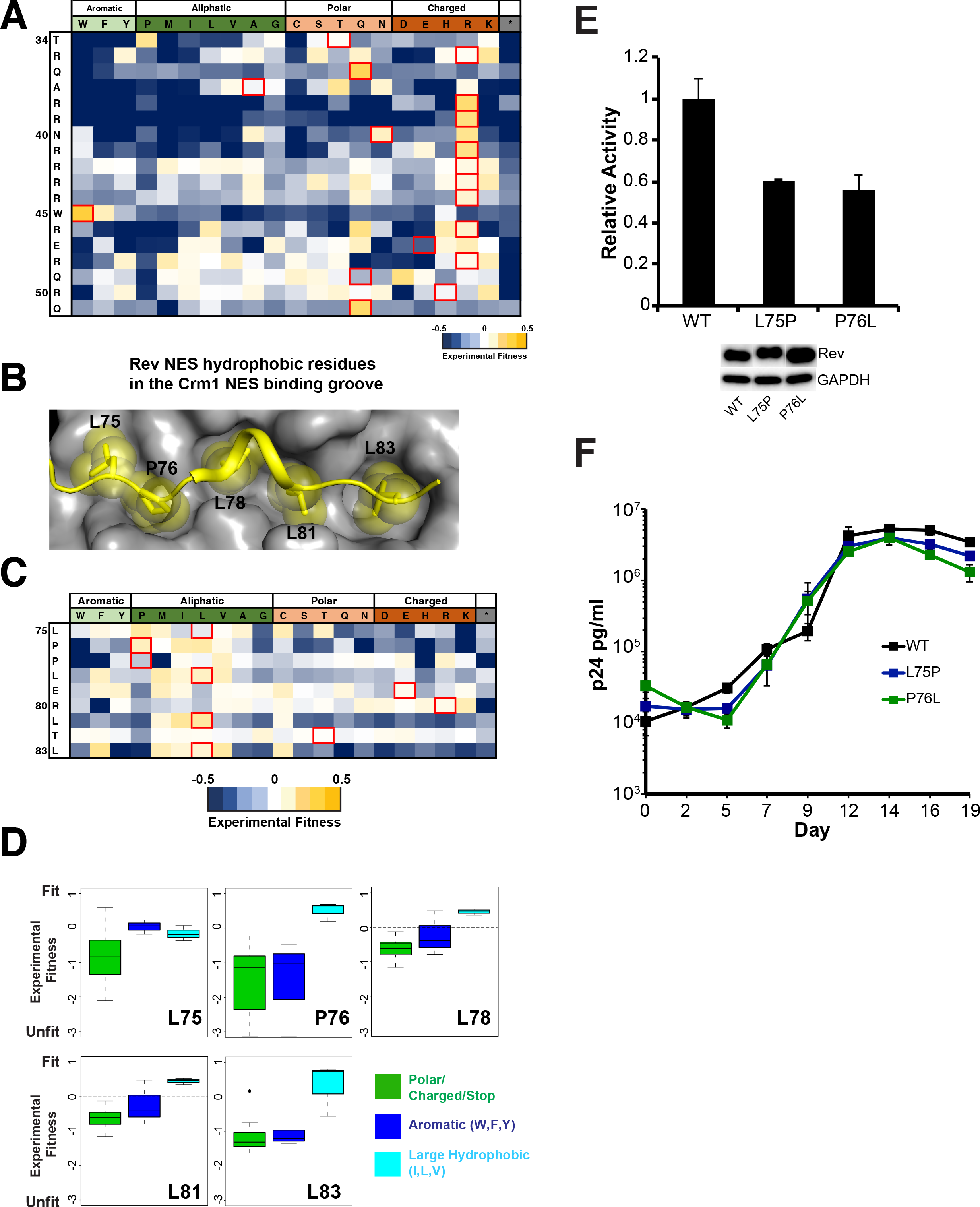
Mutational limits of the ARM and NES. A) CDMS fitness values for residues in the Rev ARM from [9]B) Structure of Rev NES (yellow) bound to Crm1 (grey) (PDBID: 3NBZ) C) CDMS fitness values for the NES from [9] D) Preferences for important Rev NES contact residues using CDMS fitness values with similar side chains grouped together [9]. E) Reporter assay probing export activity of Rev NES mutations. Data are mean ± standard deviation (s.d) of biological replicates. Western blots below show protein expression for Strep-tagged Rev and GAPDH as loading control. F) Viral replication profiles of viruses containing Rev with NES mutations. Data are mean ± s.d of biological triplicates.

The Rev NES recruits Crm1 to the Rev-RRE RNP, and recent EM studies show that a Crm1 dimer is associated with the RNP [27]. Structural studies of a Rev NES peptide bound to Crm1 have shown that Rev binds Crm1 in a non-canonical manner (Figure 3B), using prolines to properly space hydrophobic residues to contact Crm1. Curiously, CMDS data shows that P76, which contacts the Crm1 binding groove, is as fit as a hydrophobic residue (L, I), whereas L75 prefers to be a proline (Figure 3C). Furthermore, like the OD, the hydrophobic residues in the NES that contact Crm1 are tolerant of most large aliphatic side chains (Figure 3C, D).

In order to further evaluate the effect of Rev mutations within both the structured domains and linkers, we conducted a series of RNA export reporter assays together with replication assays of non-overlapped mutant viruses to avoid the confounding effects of overlapping reading frames (Figure S1) [9]. The reporter assay monitors Rev-dependent p24 capsid expression from a CMV promoter-driven truncated viral genome in 293T cells [28] and the replication assay utilizes *rev-in-nef* viruses as previously described [9]. The combination of assays also allows us to potentially uncover additional functions of Rev in the replication context.

We tested both Rev L75P and P76L mutants for function in our reporter assay and observed only 60% activity (Figure 3E), while the corresponding viral spreads were comparable to wild-type Rev, consistent with the CDMS data (Figure 3F). These observations strongly indicate that Rev is able to tolerate greater variability to mutation of Crm1 binding residues than expected and that the loss of activity observed in our reporter assay is still above some functional minimum, or that other mechanisms, present in the context of viral replication, compensate for this loss of activity.

The OD, ARM and NES are all structured, critical regions of Rev, where a point mutation is often sufficient to abolish function. While our CDMS data indicate that many residues of the OD and NES are tolerant to substitutions, conferring mutational robustness/plasticity, the ARM has more stringent sequence preferences. This may reflect the more complex nature of interactions mediated by the ARM (polar, charged, directional as well as RNA and protein), making it relatively more fragile than the OD or NES.

### The N- and C-termini influence Rev stability

While the function of the ARM, OD and NES are fairly well understood, the roles of the linkers in Rev function, apart from connecting the modules, is less clear. Since most residues within these regions do not experience strong selection for a specific amino acid/category, point mutational datasets are insufficient to determine their sequence constraints. Thus, we constructed a series of truncation and gross substitution mutations within these regions and measured their activities to interrogate their functions more rigorously.

The N-terminal 8 residues of Rev are disordered in most crystal structures (Figure 1B) [10–12, 24], but a recent structure of a scFv-bound Rev solved at 2.3 Å shows that the first helix begins at residue 5 (PDB ID: 5DHV) [24]. Genetic screens and selection assays have identified residue 12 as the first essential contact in the Rev OD [22, 23]. The presence of an overlapping reading frame encoding the critical ARM of Tat, another essential viral protein, has confounded conservation analyses of the Rev N-terminus. Indeed, previous work suggests that this region does not experience strong selective pressure (excluding requirements for translation initiation), but is well conserved in Rev in order to maintain Tat function [9].

To more fully investigate the role of N-terminus of Rev, we measured the effects of truncations in our export assay. Deleting the first 10 residues (Rev ∆1-10) had no effect on activity whereas any further truncation resulted in loss of activity (Figure 4A). Replication spreads of viruses containing these truncations recapitulated the export results, with the virus expressing Rev ∆1-10 replicating similar to the wild-type Rev virus, whereas the Rev ∆1-11 virus failed to replicate (Figure 4B). These results corroborate that the first 10 residues of Rev are not required for function.

**FIGURE 4:**
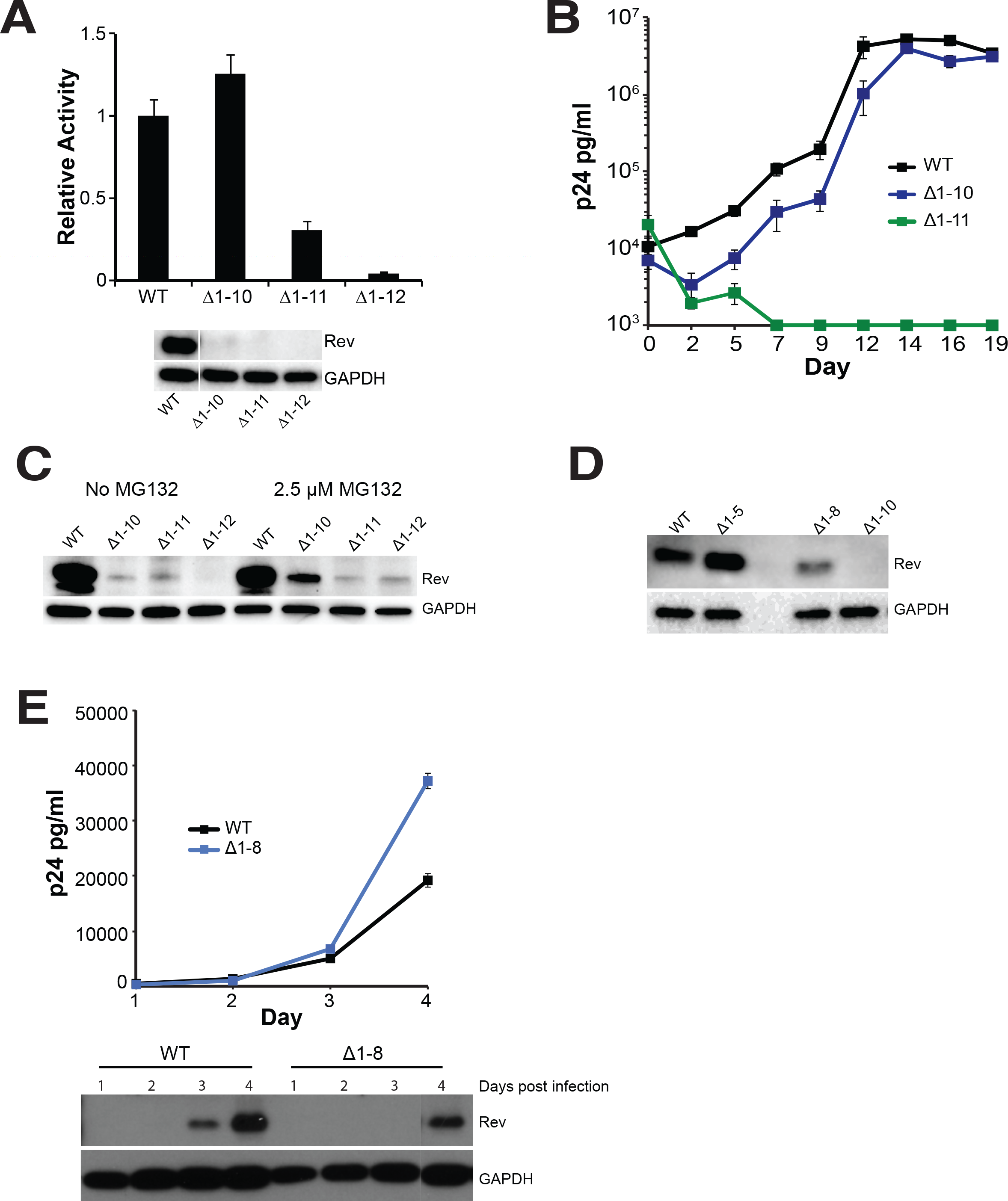
The HIV-1 Rev N-terminus is not required for function but regulates Rev stability. **A)** Reporter assay probing export activity of Rev N-terminal truncations. Data are mean ± standard deviation (s.d) of biological replicates. Western blots below show protein expression for Strep-tagged Rev and GAPDH as loading control. B) Viral replication profiles of viruses containing Rev with N-terminal truncations. Data are mean ± s.d of biological triplicates. Viral p24 levels below 1000 pg/ml are shown as 1000 pg/ml in the plots for illustration purposes. C) Adding MG132, a proteasome inhibitor partially restores expression of N-terminally truncated Rev in transiently transfected 293T cells. D) Shortening the length of N-terminal truncations restores protein expression transiently transfected 293T cells. E) N-terminal truncations affect Rev expression/stability from HIV-1 (with Flag-tagged Rev in the *nef* locus) in infected T-cells. Plot shows virus production from cell supernatant, quantified by p24 ELISA. Western blots below show protein expression for Flag-tagged Rev and GAPDH loading control from the cells. Data are mean ± s.d of biological replicates.

Unexpectedly, none of the truncated Rev proteins could be detected by western blot, despite some (such as Rev ∆1-10) being functional in both reporter and replication assays (Figure 4A, B). Previous reports have shown that Rev function is robust even at very low expression levels [29, 30]. Indeed, we observed that serial dilutions of Rev plasmid demonstrated high reporter activity at even very low levels and that activity was actually reduced at higher plasmid levels (Figure S2). Therefore, we inferred that the truncated Rev proteins were being expressed at levels sufficient for function but below the level of detection. We were able to detect low levels of truncated Rev in the presence of proteasome inhibitor, MG132 (Figure 4C), suggesting that the truncated proteins were being degraded by the proteasome. We next made smaller truncations to determine at which point Rev expression was restored and found that Rev ∆1-8 showed modest expression whereas Rev ∆1-5 showed wild-type expression. The results suggest that the truncations affect the ability of the first helix to form completely and stably (Figure 4D), thereby influencing protein stability. Remarkably, early mutational work on Rev also identified point mutations in the N-terminus (R4S5 to D4L5) that reduced Rev stability without completely abolishing function [31].

We also monitored Rev expression from NL4-3 viruses in infected T-cells, a more relevant condition. We utilized Rev with a C-terminal Flag tag (for ease of detection) in *rev-in-nef* viruses [12], infected SupT1 cells with these viruses and tracked Rev expression in cell lysates. Rev ∆1-8 displayed poorer expression than full-length Rev, confirming the reduced stability of Rev in T-cells (Figure 4E), yet conferred no defect in replication. Overall, our findings indicate that the N-terminal residues are not required for replication but nevertheless contribute to Rev stability.

The Rev C-terminus (residues 70-116) is disordered in crystal structures although recent work has suggested that it may become ordered in certain contexts. Cryo-EM studies have suggested that residues 94-116 become ordered when Rev forms filaments [24], and C-terminal truncations (1-69 and 1-93) do not form filaments under the same conditions. Evolutionary coupling analysis also suggests that the C-terminus can engage in long-range contacts and become structured [32]. Early truncation studies on Rev at various positions beyond the nuclear export sequence indicated that truncated Rev was functional [31, 33–35]. CDMS data show that selection against stop codons ends at residue 86 (the NES ends at residue 83) (Figure 5A) indicating that the C-terminus is not required for viral replication. Curiously, the distribution of fitness values for stop codons between position 86-116 is overall positive (Figure 5B; red distribution is right shifted), indicating that viruses with a truncated Rev might have a competitive advantage compared to wild-type virus and suggesting an inhibitory role for the Rev C-terminus [32]. This distribution is distinct from that of the C-terminus of Tat, another overlapped essential HIV protein, where the fitness values of stop codons appear to be near neutral after the last functional Tat domain at residue 65 (Figure 5B; blue distribution centered around zero).

**FIGURE 5:**
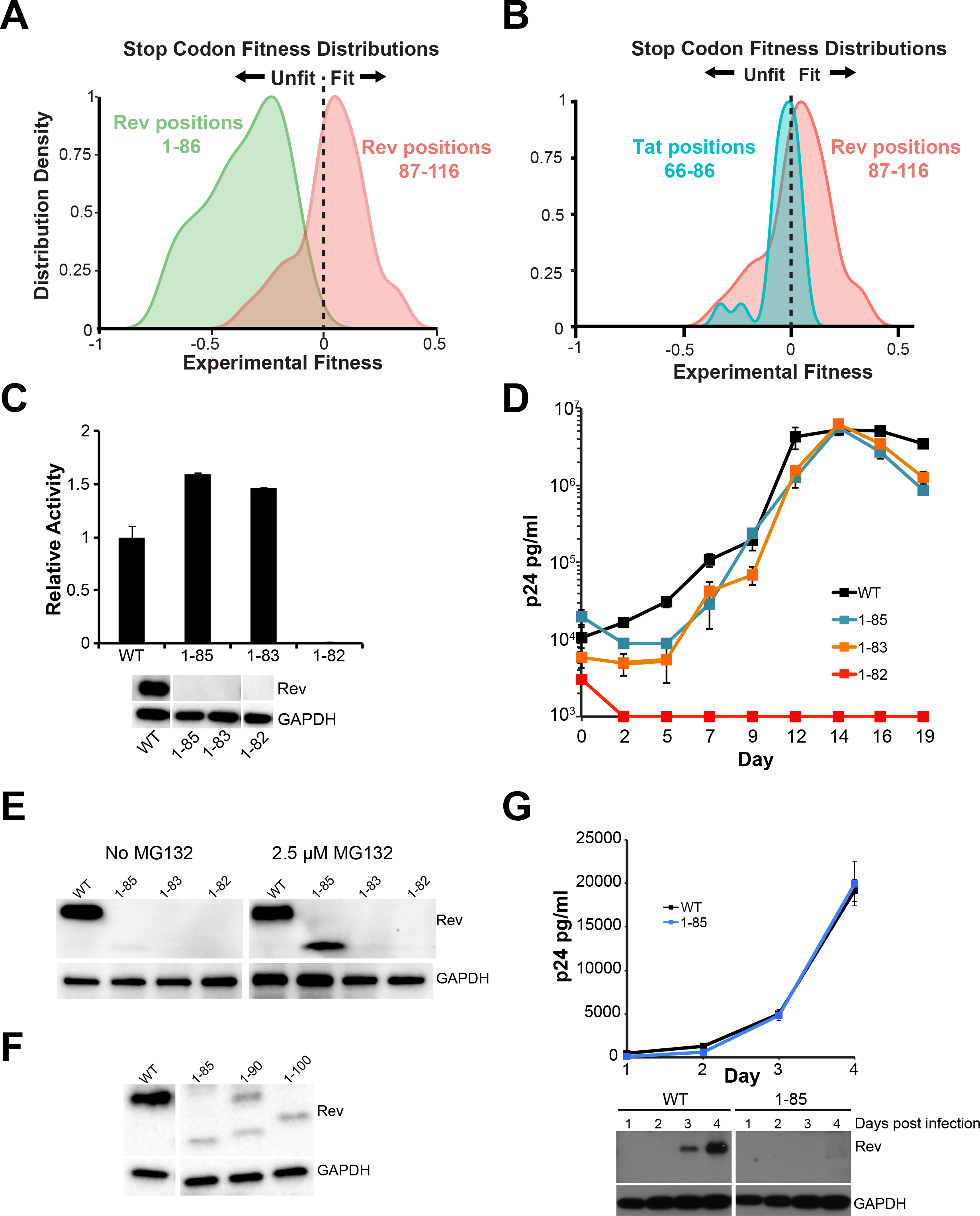
The HIV-1 Rev C-terminus is not required for function but regulates Rev stability. A) Stop codon fitness distributions from CDMS datatest for Rev [9]. B) Stop codon fitness distributions from CDMS datatest for Tat C-terminus (blue, residues 66-86) and Rev C-terminus (red, residues 86-116) (stop codon data for both proteins from [9]). C) Reporter assay probing export activity of Rev C-terminal truncations. Data are mean ± standard deviation (s.d) of biological replicates. Western blots below show protein expression for Strep-tagged Rev and GAPDH as loading control. D) Corresponding viral replication spread experiments. Viral p24 levels below 1000 pg/ml are shown as 1000 pg/ml in the plots for illustration purposes. Data are mean ± s.d of biological triplicates. E) Adding MG132, a proteasome inhibitor partially restores expression of Rev with C-terminal truncations in transiently transfected 293T cells. F) Shortening the length of C-terminal truncations does not restore protein expression transiently transfected 293T cells. Here, we note that we consistently observe two bands for Rev 1-90, one with a faster mobility and the second with the same mobility as full-length Rev. G) C-terminal truncations affect Rev expression/stability from HIV-1 (with Flag-tagged Rev in the *nef* locus) in infected T-cells. Plot shows virus production from cell supernatant, quantified by p24 ELISA. Western blots below show protein expression for Flag-tagged Rev and GAPDH loading control from the cells. Data are mean ± s.d of biological replicates.

C-terminal truncations (Rev 1-85, Rev 1-83) are functional in our reporter and replication assays, further confirming that this region is dispensable for function, while truncation of a single residue in the NES (Rev 1-82) results in a dramatic loss of function (Figure 5C, D). Similar to the N-terminal truncations, we were unable to detect Rev with C-terminal truncations despite functional reporter activity and observed low levels of truncated Rev in the presence of proteasome inhibitor, MG132 (Figure 5E). However, unlike the N-terminus, smaller truncations did not restore Rev expression as 1-85, 1-90 and 1-100 all express poorly (Figure 5F). Intriguingly, previous studies of C-terminal truncations in COS cells did not result in reduced protein stability [31], Rev 1-85 also displayed reduced stability in virus-infected T-cells, although the virus replicates as well as the wild-type Rev virus (Figure 5G). Thus, the C-terminus, like the N-terminus, is not required for RNA export or viral replication but regulates protein stability in a manner whose mechanism and functional consequence are not yet understood.

### Substitutions to the Rev turn produce different effects on RNA export and viral replication

Rev contains an 8-residue proline rich turn which separates the first two helices and spans the exon junction in the viral genome (Figure 1A, C). Recent structural studies implicate the turn residues as part of an additional Rev-Rev interaction surface where residues W45, P28, P29 and P31 “hook” with the same residues on an adjacent Rev molecule. This interface has been proposed to act as a molecular bridge between independent oligomerization events along the RRE [24, 25]. Most residues in the turn do not have strong preferences in the CDMS study (Figure 1B) [9], consistent with our understanding of the region as a simple linker. However, residue 28 demonstrated a strong preference for an aromatic side chain (W,F,Y) rather than the reference NL4-3 proline (P) (Figure S3). Interestingly, the most common allele in patient sequences is Y28, although P28 is also substantially represented and is found in many lab-adapted HIV-1 strains [36]. It is possible that a P28Y substitution could provide stacking interactions to stabilize this aromatic interface within the turn [25], but we found no significant difference in activity between the proline and tyrosine variants (Figure S3). Even an alanine substitution at position 28 is only marginally weaker in replication assays, indicating substantial sequence plasticity, but leaving the manner in which Y28 confers a fitness advantage in competition experiments and in patients still unclear. Nevertheless, given that the turn appeared mutationally plastic in our CDMS studies and since it also determines the relative orientation of the first two helices, we examined its mutational limits with respect to side chain preference, length and structure.

To this end, we created a set of mutations that replaced the turn residues (26-33) with either all alanines (turn-8A, designed to promote helical propensity [37]) or glycines and serines (turn-8GS, designed to allow greater flexibility), and monitored the effects on Rev function in both reporter and replication assays. Surprisingly, we found that both variants yielded significant levels of export in the reporter assay (Figure 6A) although neither was as active as wild-type Rev. However, shortening the length of the substitutions reduced export activity indicating that the turn must be of sufficient length to connect the two helices and produce functional Rev.

**FIGURE 6:**
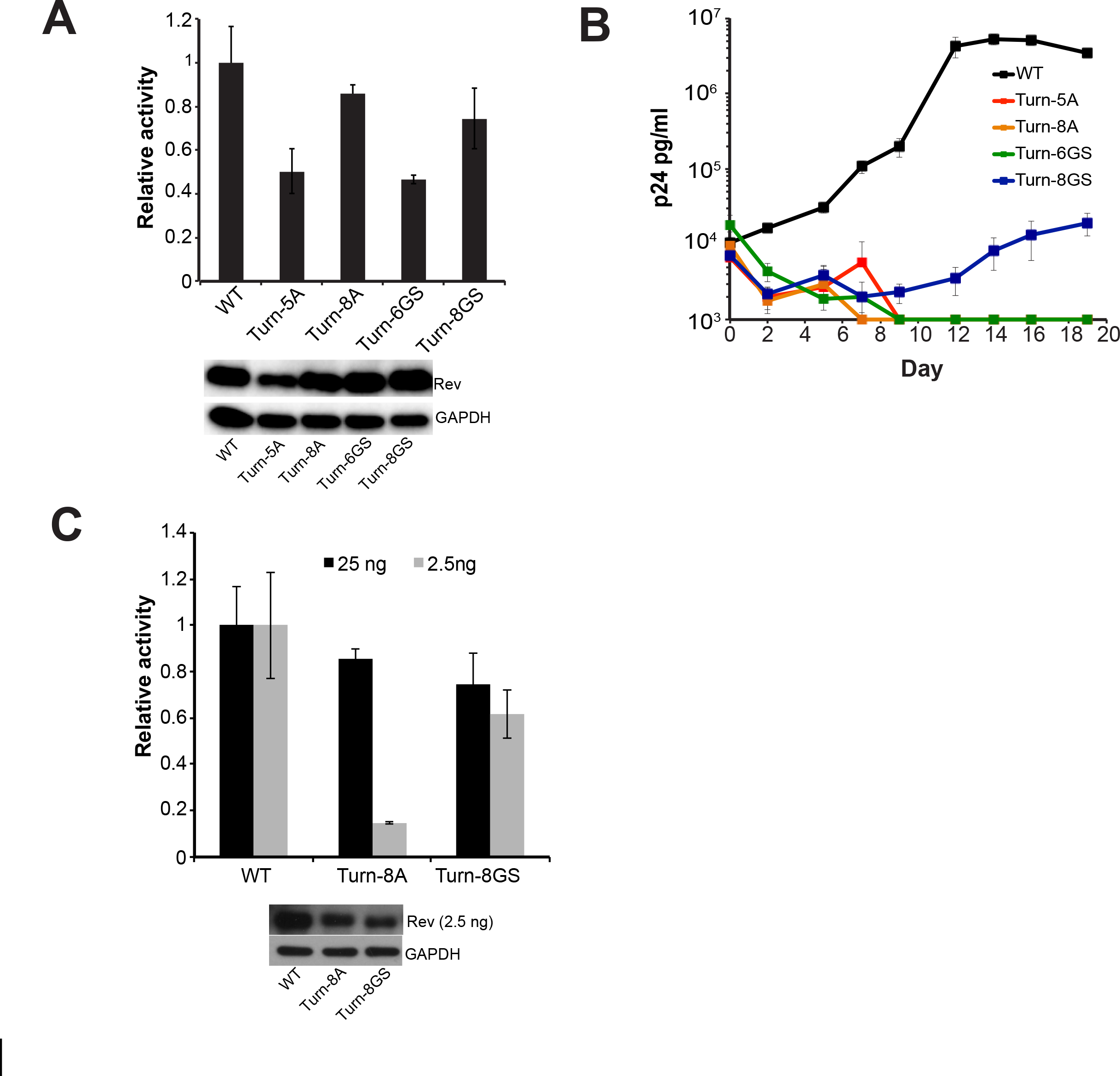
The turn plays an intriguing role in Rev function A) Reporter assay monitoring function of Rev constructs with mass substitution of turn residues 26-33. Data are mean ± standard deviation (s.d) of biological replicates. Western blots below show protein expression for Strep-tagged Rev and GAPDH as loading control. B) Corresponding viral replication spread experiments. Viral p24 levels below 1000 pg/ml are shown as 1000 pg/ml in the plots for illustration purposes. Data are mean ± s.d of biological triplicates. C) Reporter assay monitoring export activity at 25 ng and 2.5 ng of transfected Rev plasmid. Western blots below showing protein expression for Strep-tagged Rev (at 2.5 ng only) and GAPDH as loading control. Data are mean ± standard deviation (s.d) of biological replicates.

Despite significant activities in the reporter assay, all turn mutants were defective in replication, with only the turn-8GS mutant-virus showing any replication although severely attenuated (Figure 6B). Given our previous observation that Rev is functional at even low concentrations, we repeated the reporter assay with 10-fold less Rev plasmid (2.5 ng versus 25 ng) to determine if less Rev expression might recapitulate the replication results. Indeed, the export activity of the turn-8A mutant dropped significantly at low Rev expression levels (Figure 6C), indicating that it is defective and probably structurally unstable due to the replacement of a turn with a helical region. These results are consistent with previous reporter assays showing loss of function when 27PPP29 was mutated to alanines [38]. However, the turn-8GS mutant remained active at the lower Rev plasmid level unlike the turn-8A mutant (Figure 6B), despite wild-type Rev being 2-fold more active at the lower plasmid level (Figure S2). This indicates that the turn need only provide flexibility between Rev helices in order for proper export. However, the turn-8GS mutant being export competent but defective in viral replication suggests that the turn may influence Rev function in alternate pathways beyond nuclear export.

### The Rev OD-NES Linker (ONL) becomes structured

An ∼11 residue linker region (ONL) connects the second helix of Rev and the NES and is disordered in most Rev crystal structures [11, 24]. CDMS data suggests that the residues in this region can sample many amino acids with no clear preference for chemical property (Figure 1C). In order to probe the requirements of length and structure for this region, we applied a strategy similar to that used to probe the turn. We replaced residues 62-72 of the ONL with either 11 alanines (ONL-11A, to mimic a rigid, structured linker) or 11 glycine-serines (ONL-11GS, to mimic a flexible disordered linker) and tested the effect of these substitutions in reporter and replication assays. ONL-11A was nearly fully functional in the reporter assay whereas ONL-11GS showed a marked decrease in activity (Figure 7A). Viral replication spreads recapitulated this reporter data (Figure 7B), with only the structured alanine linker displaying activity. In order to test the length requirements of this region, we shortened the ONL to nine (ONL-9A) or six (ONL-6A) alanines. ONL-9A displayed reduced function in the reporter assay but was surprisingly functional for replication, whereas ONL-6A was poor in both assays (Figure 7A, B), demonstrating that the ONL has a minimum length requirement. The activity levels of the ONL-9A and ONL-11A at reduced Rev expression levels (2.5 ng of Rev plasmid) were comparable to those obtained at 25 ng of Rev plasmid, demonstrating that this behavior holds for a broad range of Rev expression levels (Figure 7C).

**FIGURE 7:**
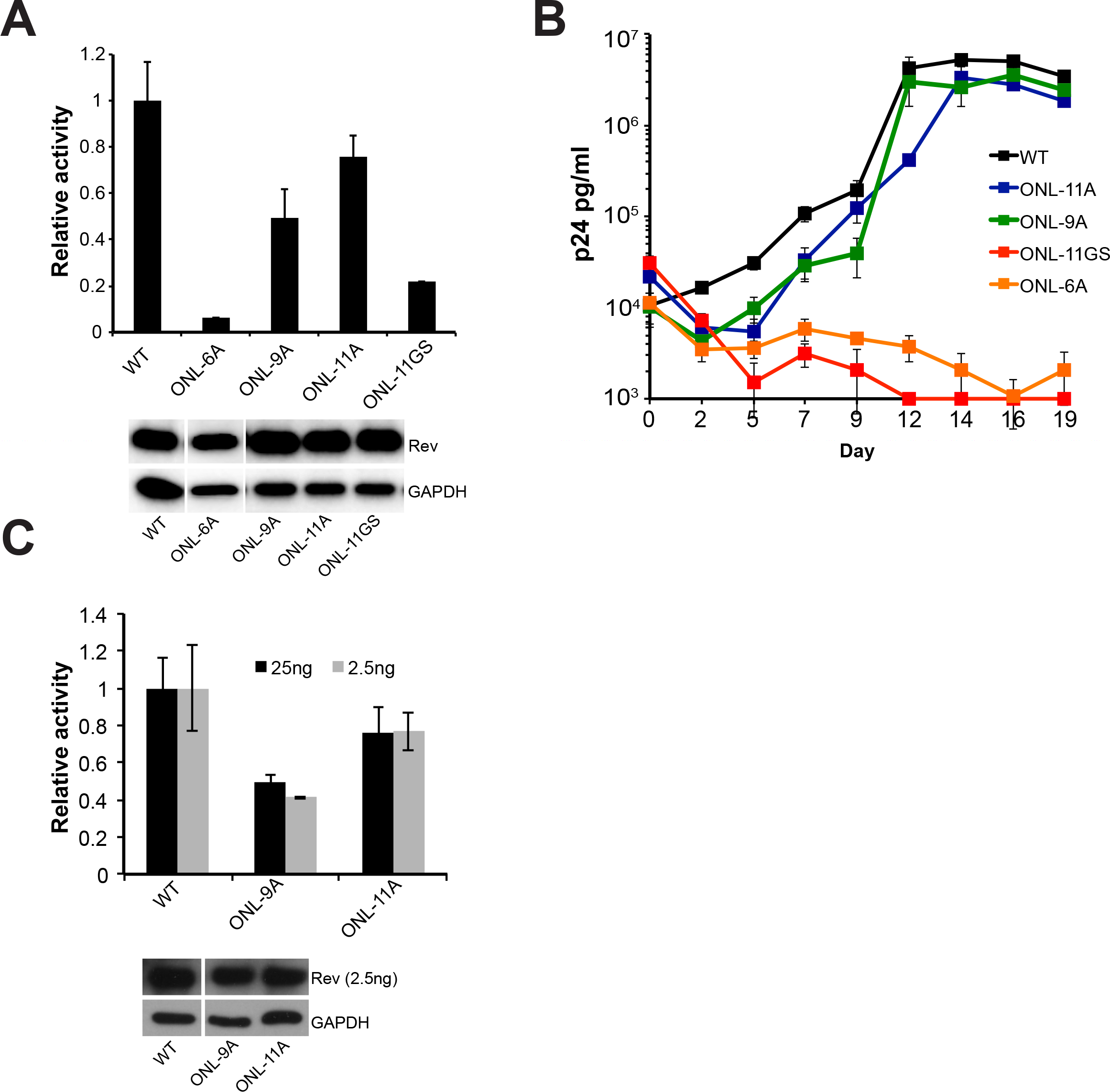
The OD-NES linker (ONL) requires structure for Rev function. A) Reporter assay monitoring export function of Rev constructs with mass substitution of ONL residues 62-72. Data are mean ± standard deviation (s.d) of biological replicates. Western blots below showing protein expression for Strep-tagged Rev and GAPDH loading control. B) Corresponding viral replication spread experiments. Viral p24 levels below 1000 pg/ml are shown as 1000 pg/ml in the plots for illustration purposes. Data are mean ± s.d of biological triplicates. D) Reporter assay monitoring export activity at 25 ng and 2.5 ng of transfected Rev plasmid. Western blots below showing protein expression for Strep-tagged Rev (at 2.5 ng only) and GAPDH as loading control. Data are mean ± standard deviation (s.d) of biological replicates.

The ability of the ONL to tolerate complete replacement with alanines but not glycine/serine suggests that this region becomes structured during Rev function, possibly during oligomeric assembly on the RRE or Crm1 binding. Our point mutational CDMS was unable to detect this requirement as it only examines the effect of individual point mutations, any one of which appears insufficient to disrupt the induced structure.

## DISCUSSION

HIV Rev has been extensively investigated over three decades. Structural studies over the past several years have led to a model in which Rev is composed of a few highly conserved, structured regions with well-defined functions that are connected by disordered linkers (Figure 1A). The results presented here expand the functional repertoire to these linker regions. Indeed, recent work in other systems has demonstrated the importance of disordered and linker regions for establishing and tuning protein interactions [39–41], and particularly for modulating RNA-protein interactions [42].

The use of disordered linkers to tune Rev function may reflect some of the selective pressures imposed by having its entire coding sequence overlapped with other HIV-1 genes, specifically *tat* and *env* (Figure 1A). In previous work, we observed a complete segregation of function within the overlapped region of *tat* and *rev*, where the overlapped nucleotides encoded amino acids critical for the function of one protein or the other, but never both. Furthermore, the genetically fragile activation domain of Tat is encoded in a non-overlapped portion of the *tat* gene, consistent with the idea that brittle domains in which many mutations are expected to be lethal are unlikely to evolve within regions that must simultaneously maintain two overlapping coding sequences. Since Rev overlaps with other genes over its entire sequence, it apparently evolved relatively mutable interaction surfaces, such as the OD, to create genetic stability. The linker regions modulate Rev function and stability yet are even more mutable and as a result are effectively segregated at the coding level, imposing few constraints on the other reading frames.

Truncations of the N- and C-termini revealed that these regions greatly influence Rev protein stability, yet have little effect on Rev function in the context of our cellular reporter and replication assays. Proteomics studies have reported similar protein interaction profiles for N-terminal Rev truncations (∆1-8) and wild-type Rev [20], indicating that N-terminus likely does not serve as a major interaction scaffold. However the N-terminus can be phosphorylated [43], and may have regulatory effects that effect the function and/or stability of the protein in patient viral replication settings.

Intriguingly, the CDMS data suggest a fitness advantage to C-terminal truncations and thus a possible inhibitory role (Figure 5A, B). A recent study proposed that the Rev NES can be masked in the cytoplasm, controlling Rev trafficking in and out of the nucleus [29]. We speculate that removing C-terminal residues might decrease the masking and thereby promote export of viral RNAs to provide a fitness advantage. Conversely, we observe that truncation of the C-terminus decreases protein stability, which might seem to negate any gains to export activity, however very little Rev is required for function, excessive amounts of Rev can lead to reduced export activity (Figure S2), and viruses harboring undetectable amounts of Rev can replicate well (Figure 4E, 5G). Hence, it appears that truncating the C-terminus might increase fitness by reducing Rev levels to a more optimal level. However, we note that our viral replication experiments are performed with *rev-in-nef*, and *nef* transcripts are nearly two-fold more abundant than *rev* transcripts [44, 45]. Therefore, some of the mutants with unaffected replication but reduced Rev stabilities might be compensated for via increased expression of Rev from the *nef* locus. Expression of truncated Rev from its endogenous locus might reveal the true limits for Rev expression and viral replication.

The Rev C-terminus length varies considerably amongst HIV-1 isolates (Figure S4). Apart from the most common 116-amino acid form of Rev, some are 123 amino acids long due to a QSQGTET insertion between residues 95 and 96; other isolates carry this insertion as well as a premature stop codon and produce a truncated, 107-amino acid Rev. Insertion of the QSQGTET sequence reduces export activity (Figure S4), further suggesting that a long C-terminus inhibits spread, although further studies will be needed to reveal the exact mechanism and functional consequences of Rev C-terminus length.

The turn between the helices of Rev is highly mutable but does have certain requirements of length and flexibility. The ability of the turn-8GS mutant to export well in the reporter assay yet fail to replicate suggests that this linker region may perform additional roles in viral replication. Moreover, structural studies have determined that the turn can form a homotypic interaction interface[24, 25], and mutations expected to stabilize this interface (P28Y) are enriched in both our CDMS data and patient data[9], yet do not have a significant effect in either reporter or replication assays. Further characterization will be required to determine the exact, and potentially novel, role of this interface and, more generally the turn itself, in Rev function.

Similarly, our manipulation of the ONL uncovered that it does not tolerate replacements that increase its propensity for disorder, suggesting that it becomes structured during some step of export complex formation. Since this stretch of residues can be replaced with alanines without significant loss of function, it appears that these residues do not form a specific interaction surface, but instead play a structural role, in which the specific sequence itself is unimportant as long as the region can adopt a presumably helical structure. Negative-stain electron microscopy has shown that a Crm1 dimer binds to the Rev-RRE complex with the two NES-binding grooves distinctly positioned to associate with two NESs from the complex [27]. A structured ONL might reduce the entropic penalty associated with Crm1 assembly and also fix NESs into a favorable orientation for this interaction, thereby improving the energetics of export complex formation.

To examine how a structured ONL might impact Rev-RRE-Crm1 complex formation, we re-visited previous models for Rev-RRE-Crm1 complex assembly [12]. Given the established plasticity in RRE structure and the hydrophobic Rev interaction surfaces, we proposed earlier that these features allow Rev-RRE complexes with varying sequences to adopt configurations favorable for Crm1 recruitment and nuclear export [12]. Adding the requirement that the OD-NES linker needs to be structured (helical with the limitations imposed by two prolines in the NL4-3 sequence), we built a model of a Rev monomer containing residues 5-83 (Figure 8A), where 83 is the last residue of the NES, from the structures of Rev (5DHV) [25] and a Rev NES-Crm1 complex (3NBZ) [14] using Chimera [46]. Previous studies established a hexamer of Rev on a minimal, functional 240 nucleotide RRE [16]. Therefore, we built three representative Rev hexamers using Rev 5-83 as described previously [12] and aligned the NES from one Rev subunit in the hexamer to a Rev-NES peptide bound to Crm1 in a model of Crm1-dimer-Rev-NES complex [27] (Figure 8B, C). Four main aspects become apparent from these models: 1) With more structured residues of Rev past its second helix, only a subset of hexamers are plausible due to steric clashes of the Rev C-terminus with other Rev subunits, the RRE or Crm1 (Figure 8C, left panel); 2) The RNA-bound Rev dimer, with its narrow crossing angle of 50° between subunits, places the two NESs in a nearly favorable configuration to associate with a Crm1 dimer (Figure 8C, left and right panels); 3) Different hexamer configurations can recruit Crm1 dimers using different combinations of Rev subunits (Figure 8C); and 4) In some hexamer configurations, more than one NES is positioned close to an NES-binding groove in one of the Crm1 subunits (Figure 8C, right panel). Given that the RRE enhances Rev binding to Crm1 [27], these configurations may in fact be the dominant class of hexamers, forming an ensemble of nuclear export complexes wherein multiple NESs may associate with Crm1 in a variety of orientations [47–49]. In this way, the structuring of the ONL may restrict the possible orientations of Rev-RRE hexamers to a subset of export-competent configurations.

**FIGURE 8:**
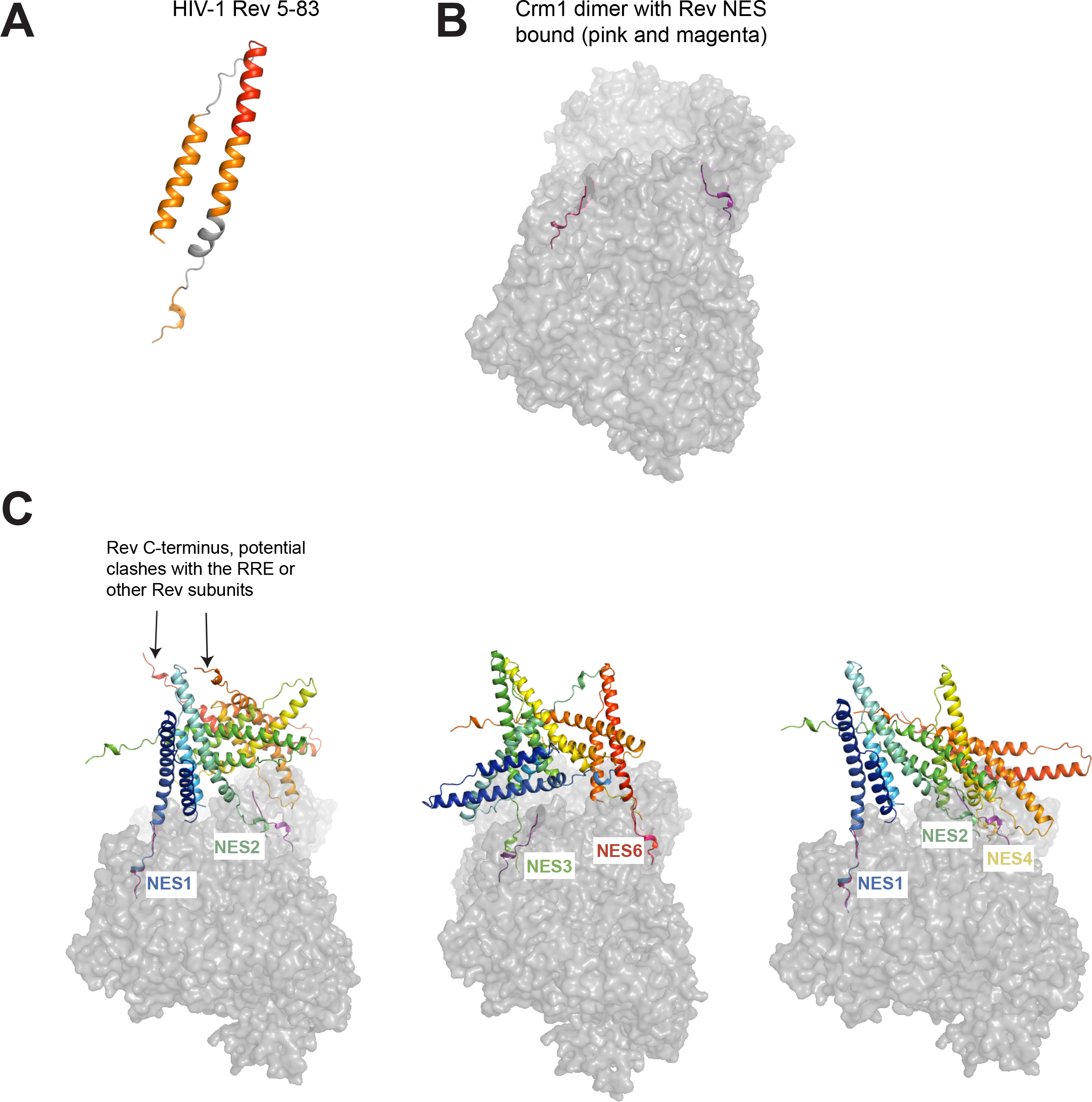
Models of HIV-1 Rev 5-83 and potential interactions of Rev hexamers with a Crm1 dimer. A) Model of HIV-1 Rev 5-83 built using Chimera B) Model of a Crm1 dimer bound to Rev NES (from [27]) C) Three representative Rev hexamers generated using Rev 5-83 and dimerization and oligomerization surfaces (as in [12]). Rev subunits are colored from blue (subunit 1) to red (subunit 6). Different Rev NES sequences were aligned with one of the Crm1-bound Rev-NES peptide: NES1 from the first panel, NES6 from the second and NES 1 from the third panel. We then observed which other Rev NES sequences were in close proximity to the second Crm1-bound Rev-NES peptide.

By our systematic analysis of each region of Rev, we conclude that every region of the protein influences its function, structure, or stability, consistent with optimal usage of limited genetic information. In the context of the viral genome and its overlapped reading frames, the virus is able to segregate essential functional and structural motifs and yet further tune Rev function using linker regions with relatively relaxed sequence and length requirements [9]. Indeed, recent analyses of Rev-like proteins in other retroviruses suggest that many key structural features - such as the key hydrophobic residues in the OD – remain conserved even as the primary sequence diverged [50]. Thus, the nuclear export function of Rev can be achieved through a wide variety of sequence configurations. Furthermore, our experiments, as well as studies of patient isolates [51], have demonstrated a wide range of Rev activity and expression levels, providing evidence of an additional levels of robustness beyond the mutational plasticity that governs protein stability and interactions studied in this work.

This study and others [52–54] show that reporter and replication assays largely correlate, suggesting that RNA export activity is the major role of Rev in the viral life cycle, however it is interesting that some Rev mutants exhibit different behaviors. For example, Rev L75P, P76L, and ONL-9A all have lowered reporter activities (40-60% of wild-type Rev) but are completely replication competent. Conversely, turn-8GS has export phenotype comparable to the above constructs but is completely defective for replication. These results illustrate that not all export defects lead to viral replication defects, suggesting that regulatory mechanisms may exist to compensate for these defects, and that reporter export activity does not always fully explain the role of Rev in the viral life cycle, raising the possibility that Rev may possess still unknown functions. Other roles for Rev, such as regulating splicing [55], translation [56–58] and encapsidation [59–61] of RRE-containing messages have been reported as well as other interactions with host partners [21, 62–65], and many of the mutants described here may help elucidate those roles.

Future work in the endogenous HIV-1 genomic context, as well as a variety of infectable cell types may shed light on these questions and reveal additional evolutionary constraints that shape Rev’s robustness. Such work will require careful design as the kinetics of Rev expression and viral spread within an organism are complex, and are regulated on multiple levels. In many of these systems, selective forces do not necessarily act to produce only a high viral titer as they do in cell culture models. The work presented here should prove valuable in deconvoluting the functional consequences of mutations in these more complex systems. In sum, our results add to an increasingly mature structural picture of Rev macromolecular complexes and demonstrate that large-scale mutagenesis approaches combined with targeted assays and structural knowledge can reveal novel and subtle mutational constraints in even well-studied systems.

## MATERIALS AND METHODS

### Construction of mutants

Primers harboring the appropriate mutations were chemically synthesized (IDT) and then used in a 2-step PCR reaction and inserted into pcDNA4TO-2xStrep-Rev (Invitrogen mammalian expression vector, described in [9]) via Gibson Assembly for testing in export assays. A similar strategy was undertaken to make viral mutants in a *rev-in-nef* virus as described previously [9, 12].

### Rev reporter assays

Reporter assays to monitor Rev export activity were carried out in HEK 293T cells (from ATCC) by transient transfection. In a typical experiment, 293T cells in a 24-well plate were transfected with 250 ng of a pCMV-GagPol-RRE reporter construct [66], 25 ng of pcDNA4TO-2xStrep-Rev (Invitrogen mammalian expression vector) (or indicated amount) and 10 ng pcDNA4TO-Firefly luciferase (to monitor transfection efficiency) using Polyjet transfection reagent (SignaGen). We chose these quantities for Rev plasmid to maintain accuracy in plasmid concentrations across different constructs and for easily detectable protein expression. Forty-eight hours after transfection, the culture supernatant was removed, the cells were lysed in Tris-buffered saline containing 1% Triton X-100 and protease inhibitors, and intracellular p24 was quantified by ELISA. Rev expression levels were verified by western blot using HRP-conjugated-anti-StrepTagII antibody (IBA biosciences) with GAPDH used as a loading control. For experiments with MG132, 2.5 uM proteasome inhibitor MG132 was added 30 hours after transfection and cells were harvested 16 hours later.

### Viral replication assays

Rev mutants were cloned into the *nef* region of a NL4-3 proviral construct, mutated to ensure no expression from the endogenous *rev* locus [12]. Virus generation and spreading assays were carried out as described earlier [9, 12]. We wish to note that the replication kinetics of an NL4-3 reference virus and an NL4-3 *rev-in-nef* virus are different, with both viruses displaying peak infection at different times (Figure S1). This is likely due differences in timing and abundance of Rev-encoding messages from an endogenous *rev* locus and a *rev-in-nef* locus.

To monitor expression of Rev-Flag during viral replication, Rev-Flag, Rev Δ1-8-Flag and Rev 1-85-Flag were cloned into the *nef* locus of a NL4-3 proviral construct and viruses were generated as above. Three million SupT1 cells (NIH AIDS Reagent Program) were infected with 150 ng of p24 (viral capsid) in a total volume of 6ml containing 1 ug/ml PEI and 8 ug/ml polybrene and centrifuged for 2 hours at 37°C at 1200xg [12]. Input virus was removed, cells were washed with 5ml of PBS and resuspended in 6 ml of media (RPMI supplemented with 25 mM HEPES pH 7.4, 10% fetal bovine serum, 1% penicillin-streptomycin. 1.25 ml of cells were withdrawn on days 1-4 post infection, lysed in RIPA buffer (20mM Tris-Cl pH 6.4, 140 mM NaCl, 1% NP-40, 0.5% sodium deoxycholate, 0.1% SDS), containing protease inhibitors and Flag-tagged Rev was detected by western blot (mouse anti-Flag antibody, Sigma) with GAPDH used as a loading control.

### CDMS analysis

Fitness values_from [9], representing selection strength based on population genetics modeling and changes in allele frequencies during viral competition assays, were used to display the preferences of each amino acid at each position. To determine each position’s general hydrophobic preferences, we grouped these fitness values into related bins (e.g. W,F,Y) and plotted their distributions.

### Patient isolate analysis

Analysis of patient sequences was performed in a manner as previously described [9]. Specifically, we downloaded high-quality, curated protein alignments from the Los Alamos HIV Sequence Compendium (http://www.hiv.lanl.gov/) and calculated amino acid conservation for every position in Rev, representing 2,694 Rev sequences. Gaps not found in the reference HXB2 sequence were discounted from the analysis.

### Analysis of stop codon fitness

Data from our CDMS experiments [9] were used to generate distributions of Rev residues 1-86 (position 86 was chosen as it is the most C-terminal position which displays selection against the stop codon), Rev residues 87-116, and Tat positions 66-86 (position 66 was chosen as it is the most N-terminal position in Tat that displays neutral or positive fitness for the stop codon). Distributions were plotted in R using ggplot2’s density function smoothed with a bandwidth adjustment parameter of 1 (adjust =1).

## ACKNOWLEDGMENTS

We thank John Gross, Arabinda Nayak, CJ Umunnakwe, Tyler Faust, and all members of the Frankel lab for helpful comments and review of the manuscript. JDF was supported in part by NIH training grant T32GM007175 and an Amgen Research Excellence Fellowship. SY was supported by the China Scholarship Council. This work was supported by NIH grant P50GM082250 to ADF.

## SUPPORTING INFORMATION CAPTIONS

**FIGURE S1:**
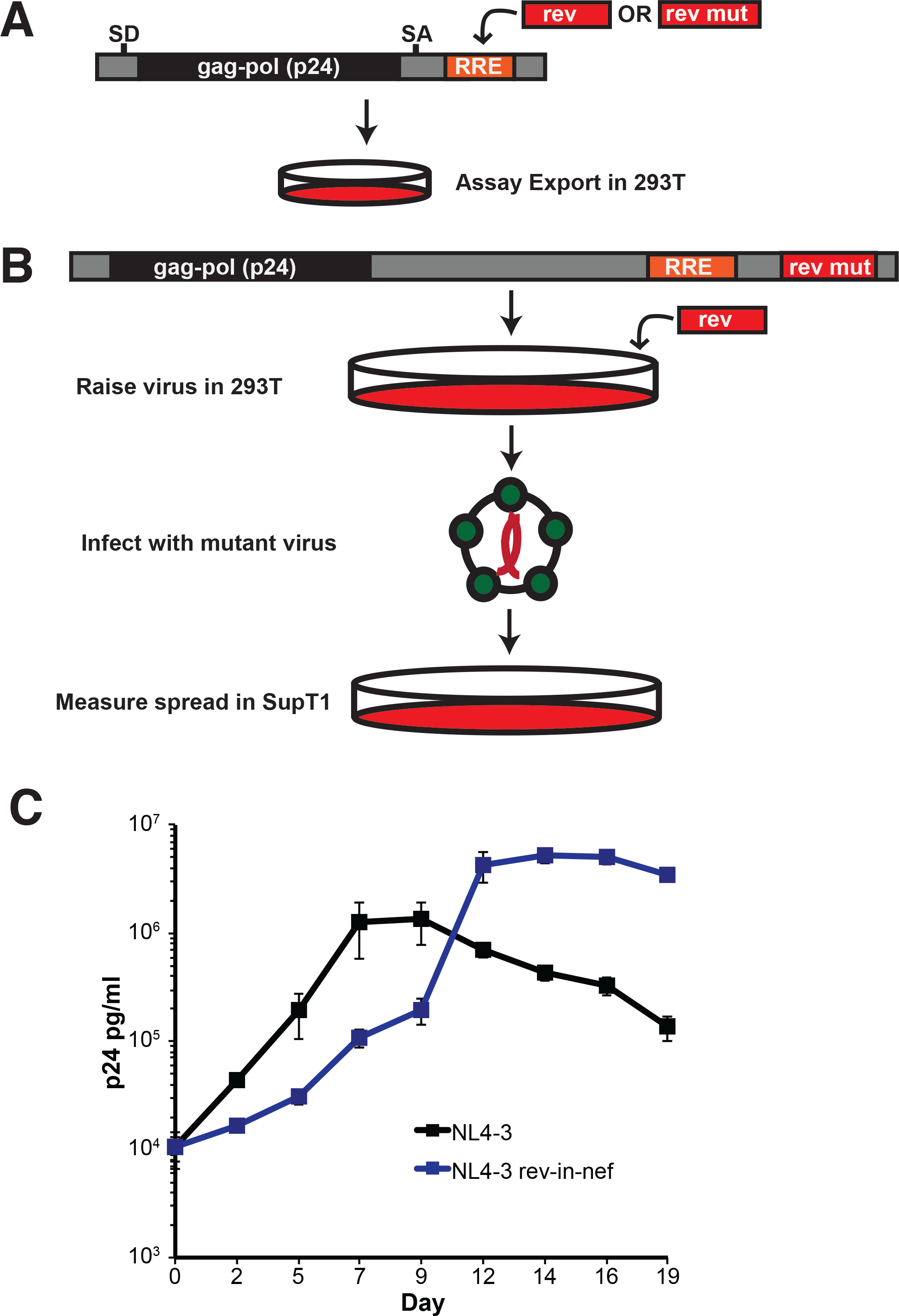
Rev functional assays. A) Reporter Assay: The Gag-Pol proteins are encoded in an intron between a splice donor (SD) and splice acceptor (SA); Rev or a Rev mutant is provided in trans and co-expression of these two constructs results in production of the Gag-Pol product, capsid protein p24 which is measured by ELISA. B) Non-overlapped viral Replication Assay. A provirus where the endogenous locus of *rev* is ablated and a *rev* mutant is inserted into the *nef* locus is used to produce viral particles in 293T cells (with wild-type *rev* provided in trans to ensure viral particle production). The mutant virus is then used to infect SupT1 cells and viral spread is measured by p24 ELISA. C) Viral replication spread assays comparing NL4-3 *rev-in-nef* virus to the NL4-3 virus.

**FIGURE S2:**
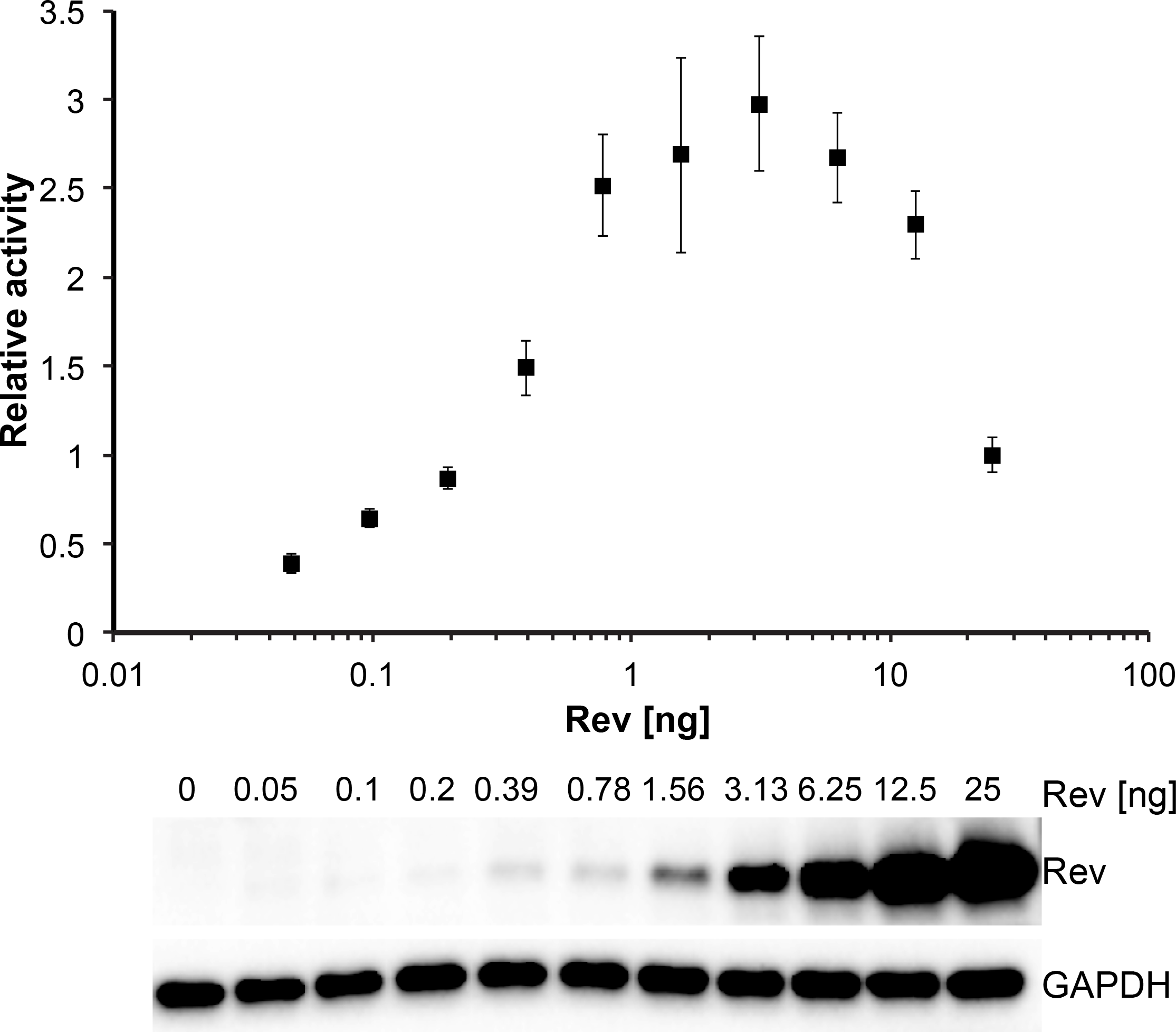
Reporter activity for a serial 2-fold dilution of transfected Rev plasmid with constant amount of reporter plasmid. Data are mean ± standard deviation (s.d) of biological replicates. Western blots below showing protein expression for Strep-tagged Rev and GAPDH loading control.

**FIGURE S3:**
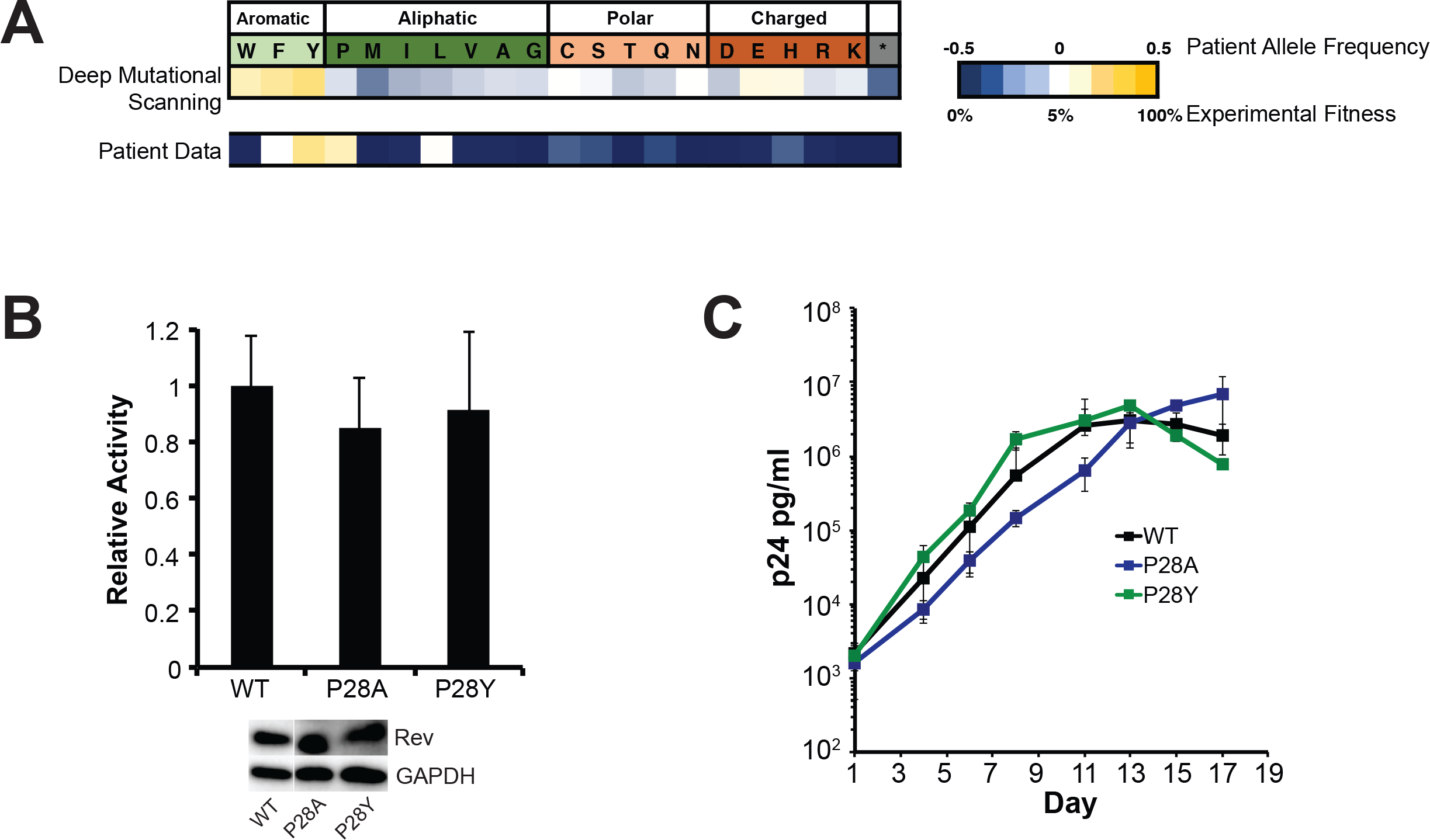
Residue 28 in the Turn A) Experimental fitness of residues for position 28 from the CDMS data B) Reporter assay monitoring export activity of mutations at residue 28. Western blots below showing protein expression for Strep-tagged Rev and GAPDH loading control. C) Corresponding viral replication spread experiments. Data are mean ± s.d of biological triplicates.

**FIGURE S4:**
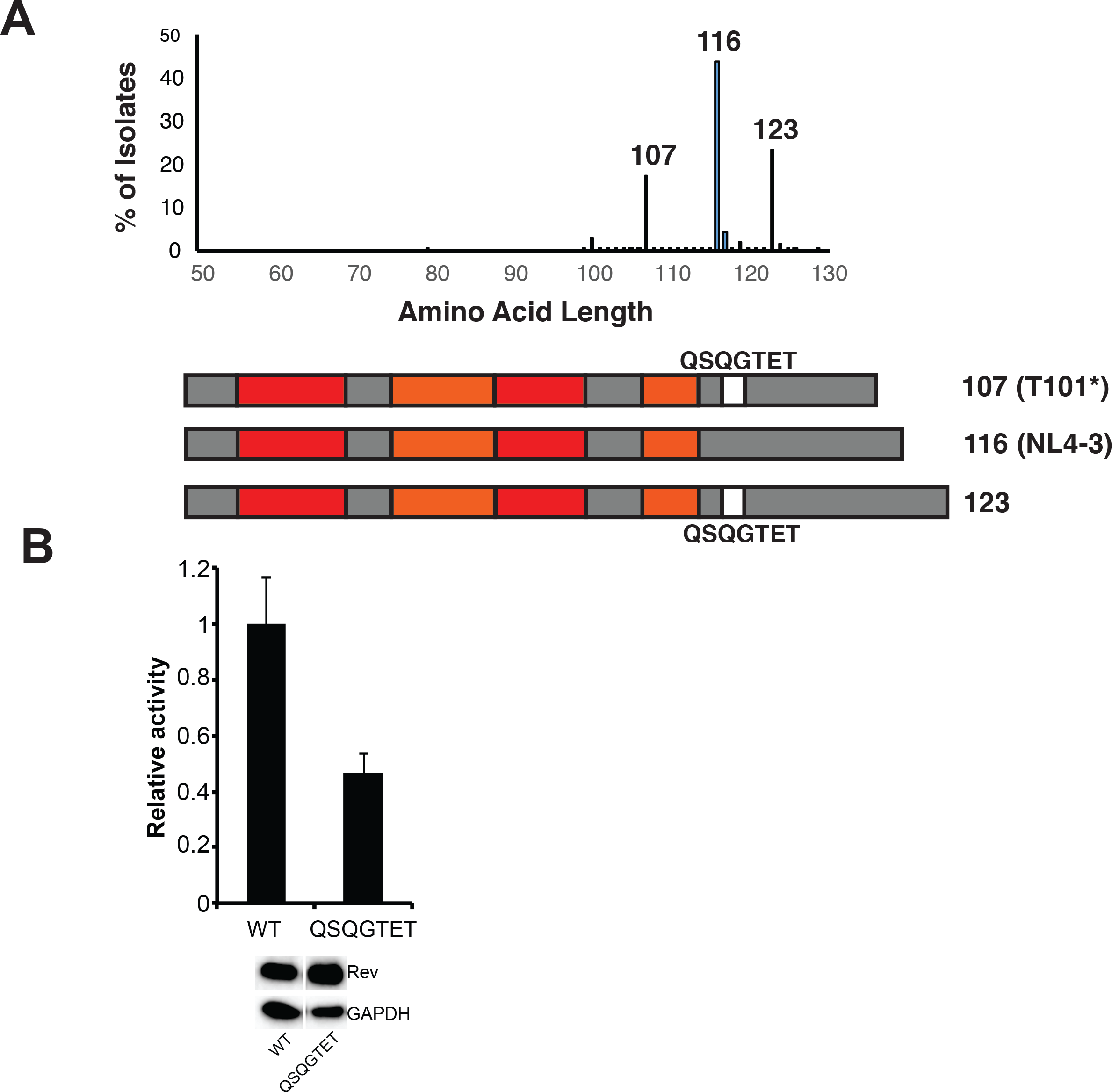
A) Length distributions of common Rev isolates observed in patients and domain organizations of major isolates with coloring as in Figure 1 and numbering according to HXB2 reference sequence. B) Reporter assay monitoring export activity of a ‘QSQGTET’ insertion into the Rev C-terminus. Western blots below showing protein expression for Strep-tagged Rev and GAPDH loading control.

